# Multivariate pattern connectivity

**DOI:** 10.1101/046151

**Authors:** Stefano Anzellotti, Alfonso Caramazza, Rebecca Saxe

## Abstract

When we perform a cognitive task, multiple brain regions are engaged. Understanding how these regions interact is a fundamental step to uncover the neural bases of behavior. Most research on the interactions between brain regions has focused on the univariate responses in the regions. However, fine grained patterns of response encode important information, as shown by multivariate pattern analysis. In the present article, we introduce and apply multivariate pattern connectivity (MVPC): a technique to study the dependence between brain regions in humans in terms of the multivariate relations between their patterns of responses. MVPC characterizes the responses in each brain region as trajectories in region-specific multidimensional spaces, and models the multivariate relationship between these trajectories. Considering the fusiform face area (FFA) as a seed region, we used searchlight-based MVPC to reveal interactions between regions undetected by univariate functional connectivity analyses. MVPC (but not functional connectivity) identified significant interactions between right FFA and the right anterior temporal lobe, the right superior temporal sulcus, and the dorsal visual stream. Additionally, MVPC outperformed univariate connectivity in its ability to explain independent variance in the responses of individual voxels. In the end, MVPC uncovered different connectivity profiles associated with different representational subspaces of FFA: the first principal component of FFA shows differential connectivity with occipital and parietal regions implicated in the processing of low-level properties of faces, while the second and third components show differential connectivity with anterior temporal regions implicated in the processing of invariant representations of face identity.

**Author Summary:** Human behavior is supported by systems of brain regions that exchange information to complete a task. This exchange of information between brain regions leads to statistical relationships between their responses over time. Most likely, these relationships do not link only the mean responses in two brain regions, but also their finer spatial patterns. Analyzing finer response patterns has been a key advance in the study of responses within individual regions, and can be leveraged to study between-region interactions. To capture the overall statistical relationship between two brain regions, we need to describe each region’s responses with respect to dimensions that best account for the variation in that region over time. These dimensions can be different from region to region. We introduce an approach in which each region’s responses are characterized in terms of region-specific dimensions that best account for its responses, and the relationships between regions are modeled with multivariate linear models. We demonstrate that this approach provides a better account of the data as compared to standard functional connectivity, and we use it to discover multiple dimensions within the fusiform face area that have different connectivity profiles with the rest of the brain.

## 1 Introduction

Cognitive tasks recruit multiple brain regions (Ishai, 2008; Anzellotti and Caramazza, 2015; Gallagher and Frith, 2003; Fedorenko and Thompson-Schill, 2014). How do these regions work together to generate behavior? A variety of methods have been developed to study connectivity both in terms of the anatomical structure of the brain (Le Bihan et al., 2001), and of the relations between timecourses of responses during rest (Biswal et al., 1995) and during specific experimental tasks (Friston et al., 2003; Roebroeck et al., 2005; Baldassano et al., 2012; Hermundstad et al., 2013). Functional Magnetic Resonance Imaging (fMRI) has proven to be a valuable instrument in this enterprise, offering noninvasive recording with good spatial resolution and whole-brain coverage.

In parallel to this literature, multivariate pattern analysis (MVPA; Haxby et al. (2001)) has drastically increased the potential of fMRI for the investigation of representational content, making it possible to detect information at a level of specificity that was unthinkable with previous univariate analyses (Kriegeskorte et al., 2007; Nestor et al., 2011; Anzellotti et al., 2013; Soon et al., 2008; Koster-Hale et al., 2013). Despite the success of MVPA, relatively few attempts have been made to transport the potential of multivariate analyses to the domain of dynamics and connectivity.

A recent study (Coutanche and Thompson-Schill, 2014) used trial-by-trial classification accuracy of color and shape in area V4 and in the lateral occipital complex (LOC) to predict trial-by-trial accuracy of object classification in the anterior temporal lobe (ATL). Earlier work by the same group (Coutanche and Thompson-Schill, 2013) used a continuous measure of classification based on correlations, offering a richer description of each brain region’s patterns. These studies are important steps towards exploiting the wealth of information encoded in patterns of BOLD response to study connectivity, but they both characterize the information encoded in a brain region using a single measure (a given classification), rather than in terms of values along multiple dimensions.

An additional property of both these methods (Coutanche and Thompson-Schill, 2013, 2014) is that they use classification along experimenter defined categories. This approach can be useful to probe a specific hypothesis about a given classification. However, it might disregard other information encoded by the regions studied which is orthogonal to the categories chosen by the experimenter. As a consequence, the results depend on the experimenter’s choice of the categories, and on how well the chosen categories capture the functional role of the regions studied.

Multivariate pattern connectivity (MVPC) is a novel method to investigate the connectivity between brain regions in terms of multivariate spatial patterns of responses. In keeping with the statistical literature (Mandelbrot and Wallis, 1969), we will use the term ‘statistical dependence’, which we consider more accurate. The method is composed of three main stages. In the first stage, the representational space in each brain region is modeled extracting a set of data-driven dimensions (rather than chosen by the experimenter), that correspond to spatial response patterns that ‘best’ characterize that region’s responses over time. In the second stage, the multivariate timecourses of responses in each region are reparametrized as trajectories in the representational spaces defined by these dimensions. In the third stage, the multivariate relations between the trajectories in the representational spaces of different regions are modelled. In a procedure analogous to MVPA, independent data are used to train and test the models. The dimensions and the parameters modelling the relationship between two regions are estimated with all runs but one, and then used to model the relation between those regions in the remaining run.

In a set of analyses investigating the connectivity between the fusiform face area and the rest of the brain (with a searchlight approach), MVPC identified dependencies between regions not detected by standard functional connectivity, and explained more variance in individual voxels responses than univariate methods. In the end, MVPC revealed different connectivity profiles associated with different dimensions of FFA’s responses.

## 2 Materials and Methods

### 2.1 Ethics statement

The volunteers’ consent was obtained according to the Declaration of Helsinki (BMJ, 1991, pp. 302, 1194). The project was approved by the Human Subjects Committees at the University of Trento and Harvard University.

### 2.2 Participants

A total of ten volunteers (N = 3 female, age range 18-50, mean 27.1) participated in the experiment. Data from one participant were discarded from the analysis because of poor performance during a behavioral training session administered on the day before the scanning.

### 2.3 Stimuli

Computer generated 3D models (using DAZ-3D) of 5 face identities were used to generate images at 5 different orientations for each identity (Figure 1). Stimuli were presented with Psychtoolbox (Brainard, 1997; Pelli, 1997) running on MATLAB, with the add-on ASF (Schwarzbach, 2011), using an Epson EMP 9000 projector. Images were projected on a frosted screen at the top of the bore, viewed through a mirror attached to the head coil.

**Figure 1:**
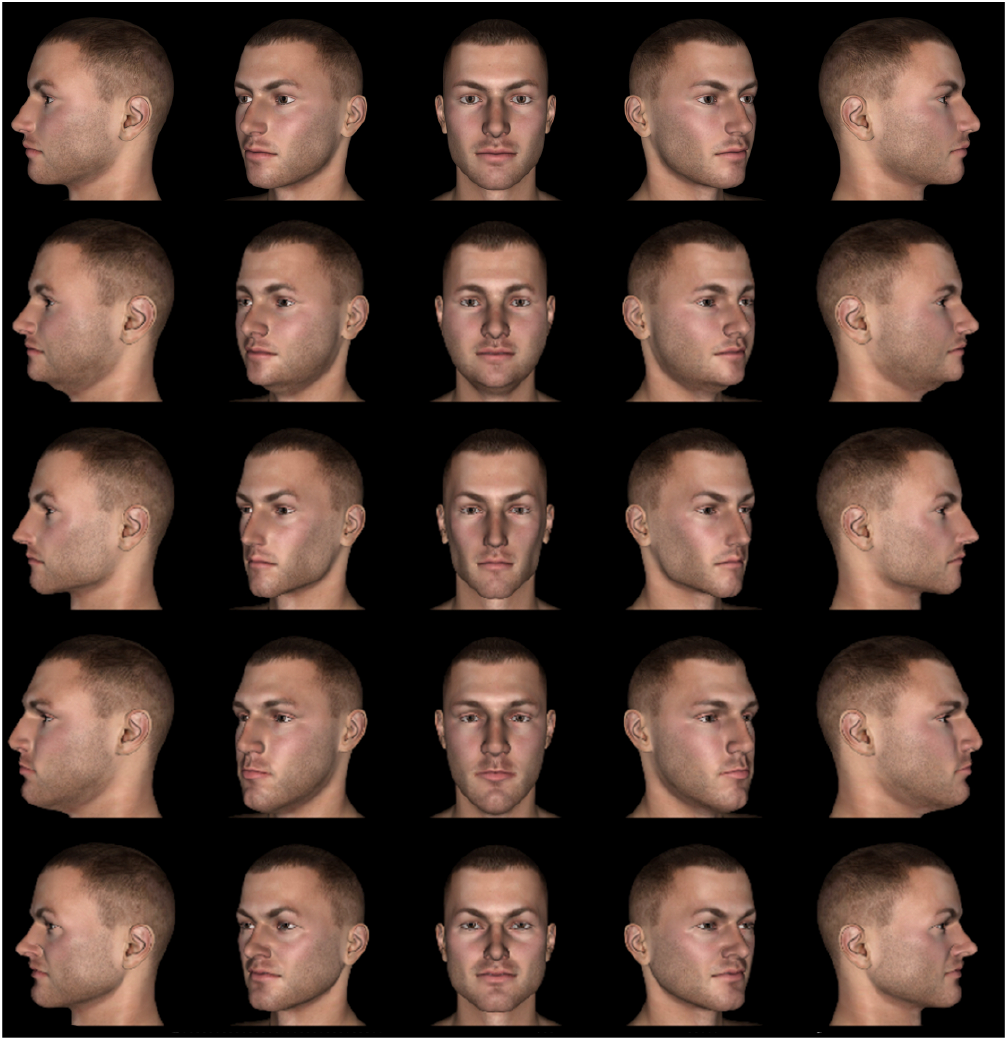
Experimental stimuli.

### 2.4 Experimental design

One of the five face identities was designated as the target, and participants were instructed to respond with the index finger of the right hand to the target face and with the middle finger to the other distractor faces (Figure 1A). Each trial consisted of the presentation of a face image (500ms) followed by a fixation cross (1500ms). The experiment was composed of three 12-minute runs, each consisting of approximately 320 trials. The order of presentation of the stimuli was generated with optseq2 (http://surfer.nmr.mgh.harvard.edu/optseq/). A 6 minutes block-design functional localizer was administered at the beginning of the fMRI session. Participants observed 16 second long blocks comprising 8 images of faces, 8 images of houses, or 8 scrambled images, and performed a 1-back task in which they had to detect repetitions of identical stimuli. None of the faces shown in the localizer were presented during the other parts of the experiment.

### 2.5 Data acquisition

The data were collected on a Bruker BioSpin MedSpec 4T at the Center for Mind/Brain Sciences (CIMeC) of the University of Trento using a USA Instruments eight-channel phased-array head coil. Before collecting functional data, a high-resolution (1 × 1 × 1 *mm*^3^) T1-weighted MPRAGE sequence was performed (sagittal slice orientation, centric phase encoding, image matrix = 256 × 224 [Read × Phase], field of view = 256 × 224 *mm*^2^ [Read×Phase], 176 partitions with 1 *mm* thickness, GRAPPA acquisition with acceleration factor = 2, duration = 5.36 minutes, repetition time = 2700, echo time = 4.18, TI = 1020 *msec*, 7° flip angle). Functional data were collected using an echo-planar 2D imaging sequence with phase oversampling (image matrix = 7064, repetition time = 2000 *msec*, echo time = 21 *msec*, flip angle = 76°, slice thickness = 2 *mm*, gap = 0.30 *mm*, with 3 × 3 *mm* in plane resolution). Over three runs, 1095 volumes of 43 slices were acquired in the axial plane aligned along the long axis of the temporal lobe.

### 2.6 Preprocessing and De-noising

Data were preprocessed with SPM12 (http://www.fil.ion.ucl.ac.uk/spm/software/spm8/) and regions of interest were generated with MARSBAR (Brett et al., 2002) running on MATLAB 2010a. Subsequent analyses were performed with custom MATLAB software. The first 4 volumes of each run were discarded and all images were corrected for head movement. Slice-acquisition delays were corrected using the middle slice as reference. Images were normalized to the standard SPM12 EPI template and resampled to a 2 *mm* isotropic voxel size. The BOLD signal was high pass filtered at 128s and prewhitened using an autoregressive model AR(1). Outliers were identified with the artifact removal tool (ART), using both the global signal and composite motion as recommended in. Datapoints exceeding experimenter-defined thresholds were removed from the analysis. An additional noise-removal step was performed with CompCor (Behzadi et al., 2007). In each individual participant, a control region was defined combining the white matter and cerebrospinal fluid masks obtained with SPM segmentation, and five principal components were extracted. Since the control region does not contain gray matter, its responses are thought to reflect noise. For each run, the timecourses of the components extracted from the control region were regressed out from the timecourses of every voxel in gray matter.

### 2.7 Searchlight

For each participant, we defined a seed region of interest in the right fusiform face area (FFA) using the independent functional localizer. Data were modeled with a standard GLM using SPM12, and the seed ROI was defined in each individual participant as a 6mm radius sphere centered in the FFA peak for the faces vs houses contrast.

We defined a gray matter mask by smoothing (with a 6mm FWHM gaussian kernel) and averaging the gray matter probabilistic maps obtained during segmentation. The average maps were then thresholded obtaining approximately 130000 gray matter voxels (127821). For each voxel in the gray matter mask, we defined a 6mm radius sphere centered in that voxel, and calculated the statistical dependence between FFA responses and responses in the sphere.

### 2.8 Standard functional connectivity

Functional connectivity was calculated low-pass filtering at 0.1 *Hz* the mean response in the seed region and the mean response in the searchlight spheres, and calculating Pearson’s correlation between the low-pass filtered responses in the seed and each sphere, thus obtaining a whole-brain functional connectivity map. Statistical significance across participants was assessed with statistical nonparametric mapping (Nichols and Holmes, 2002) using the SnPM extension for SPM (http://warwick.ac.uk/snpm).

### 2.9 MVPC: modelling representational spaces

Let us consider the multivariate timecourses *X_se,1_*,…, *X_se,m_* and *X_sp,1_*,…, *X_sp,m_* in the seed region and a sphere respectively, for experimental runs from 1 to *m*. Each multivariate timecourse *X_se,i_* is a matrix of size *T_i_* × *n_se_*, where *n_se_* is the number of voxels in the seed region and *T_i_* is the number of timepoints in run *i*. Analogously, each multivariate timecourse *X_sp,i_* is a matrix of size *T_i_ × n_sp_*, where *n_sp_* is the number of voxels in the sphere. Data analysis followed a leave-one-run-out procedure: for each choice of an experimental run *i*, data in the remaining runs were concatenated, obtaining

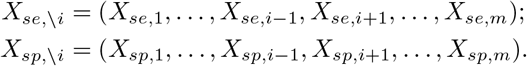

Principal component analysis (PCA) was applied to *X_se,\i_* and *X_sp,\i_*:

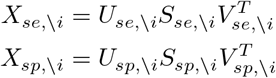

Dimensionality reduction was implemented projecting *X_se,\i_* and *X_sp,\i_* on lower dimensional subspaces spanned by the first *k_se_* and *k_sp_* principal components respectively:

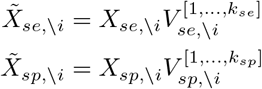

where 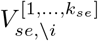 is the matrix formed by the first *k_se_* columns of *V_se,\i_* and 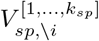 is the matrix formed by the first *k_sp_* columns of *V_sp,\i_*. In the first analysis, the number of components *k_se_* and *k_sp_* was chosen for each sphere and iteration using the Bayesian Information Criterion (BIC). In the second analysis, the incremental contribution of each component was tested by comparing the results obtained choosing 1, 2 and 3 components. We can take a moment to reflect on the interpretation of the procedure we just completed. For each region, each dimension obtained with PCA is a linear combination of the voxels in the region, whose weights define a multivariate pattern of response over voxels. Considering as an example the seed region, the loadings of a dimension *j* are encoded in the *j*-th column of 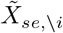, and represent the intensity with which the multivariate pattern corresponding to dimension *j* is activated over time.

### 2.10 MVPC: modelling statistical dependence

The mapping from the dimensionality-reduced timecourses in the sphere 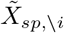 to the dimensionality-reduced timecourses in the seed 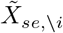 was modeled with multiple linear regression:

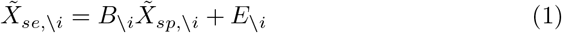

the model parameters were estimated using ordinary least squares (OLS).

### 2.11 MVPC: predicting multivariate timecourses

After having estimated parameters *B_\i_*, predictions for the multivariate responses in the left out run *i* were generated by 1) projecting the sphere data in run *i* on the sphere dimensions estimated with the other runs, and 2) multiplying them by the parameters estimated using data from the other runs. More formally, for each run *i*, we generated dimensionality reduced responses in the sphere:

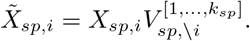

Then, we calculated the predicted responses in the seed region in run *i*:

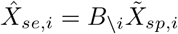

using the parameters *B_\i_*, independently estimated with the other runs.

In keeping with the use of correlation in standard functional connectivity, we calculated the correlation between the predicted and observed timecourses in each dimension in the seed region. First, we projected the observed voxel-wise timecourses in the seed region onto the lower dimensional subspace using 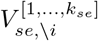:

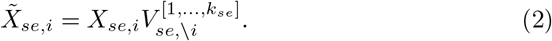

Then, we computed

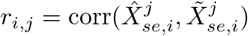

for each dimension *j* = 1,…, *k_se_* of the seed region’s subspace. In the end, we generated a single summary measure 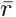, computing the average of the values *r_j_* weighted by the proportion of variance explained by the corresponding dimensions *j*:

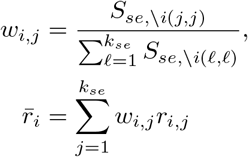

(see the relationship between the eigenvalues along the diagonal of *S* and variance explain in PCA). The values 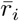 obtained for the different runs were averaged yielding 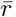. This procedure was repeated for each searchlight sphere, obtaining a whole brain map of 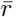 values for each participant. The significance of 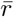 was tested across participants with statistical nonparametric mapping (Nichols and Holmes, 2002) using the SnPM extension for SPM (http://warwick.ac.uk/snpm).

### 2.12 Voxelwise variance explained

The value 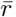 is a convenient measure of statistical dependence: it reflects how well the prediction generated by MVPC correlates with the observed data. However, in this measure, the target for prediction is the multivariate timecourse 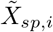. Instead, ‘standard’ univariate connectivity based on the mean timecourse aims to predict a different target: mean(*X_se,i_*, 2). Since the targets of prediction are different, the comparison between correlation obtained with mean-based univariate connectivity and the 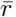 value in MVPC is less than ideal. Specifically, the correlation obtained with mean-based univariate connectivity ‘gives up’ on predicting variability orthogonal to the mean: failures to predict variability orthogonal to the mean do not lower the correlation value. In contrast, the 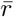 value can be penalized for poor predictions of dimensions orthogonal to the mean. As a consequence, MVPC might predict the mean response as well as mean-based univariate connectivity and still receive a lower 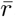 value due to poorer predictions along other dimensions. To make the predictions generated by univariate connectivity based on the average timecourse and MVPC more comparable, we need a measure of their ability to predict a common target. For this reason, for each searchlight sphere we calculated the average variance explained by mean-based univariate connectivity and MVPC in the timecourses of individual voxels in the seed region. To calculate variance explained by both methods in independent data, we needed to use a variant of functional connectivity that can perform leave-one-out predictions. The variance explained in functional connectivity is *r*^2^, and it is equal to the variance explained by a linear regression estimated and tested in the same data. We used linear regression estimated in all runs minus one, and tested the variance explained in the left-out run, thus obtaining a leave-one-out variant of mean-based univariate functional connectivity (that uses the same data-split used in MVPC). The linear regression yielded a prediction of the mean response in the seed region. Each voxel’s response was then predicted with the predicted mean response in the seed region. For MVPC, we predicted each voxel’s response projecting the multivariate prediction 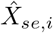 from its low-dimensional subspace of principal components to voxel space, using the matrix 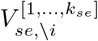. Each voxel’s response was reconstructed as the sum of the dimensions’ loadings on the voxels weigthed by the dimensions’ loadings at each timepoint. It can be helpful here to note that this is equivalent to the product

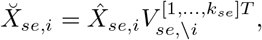

where 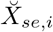 is the voxel-wise prediction (see 2 and consider that 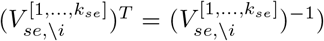. In the case of the mean-based univariate functional connectivity, the voxelwise prediction can be written as

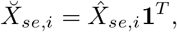

where 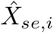 is the predicted mean response in the seed region and **1** is a *n_se_ × T_i_* vector of ones, making explicit the common form of the prediction for MVPC and for mean-based univariate connectivity: in the latter the mean is treated as a single dimension with equal loadings for each voxel.

**Figure 2:**
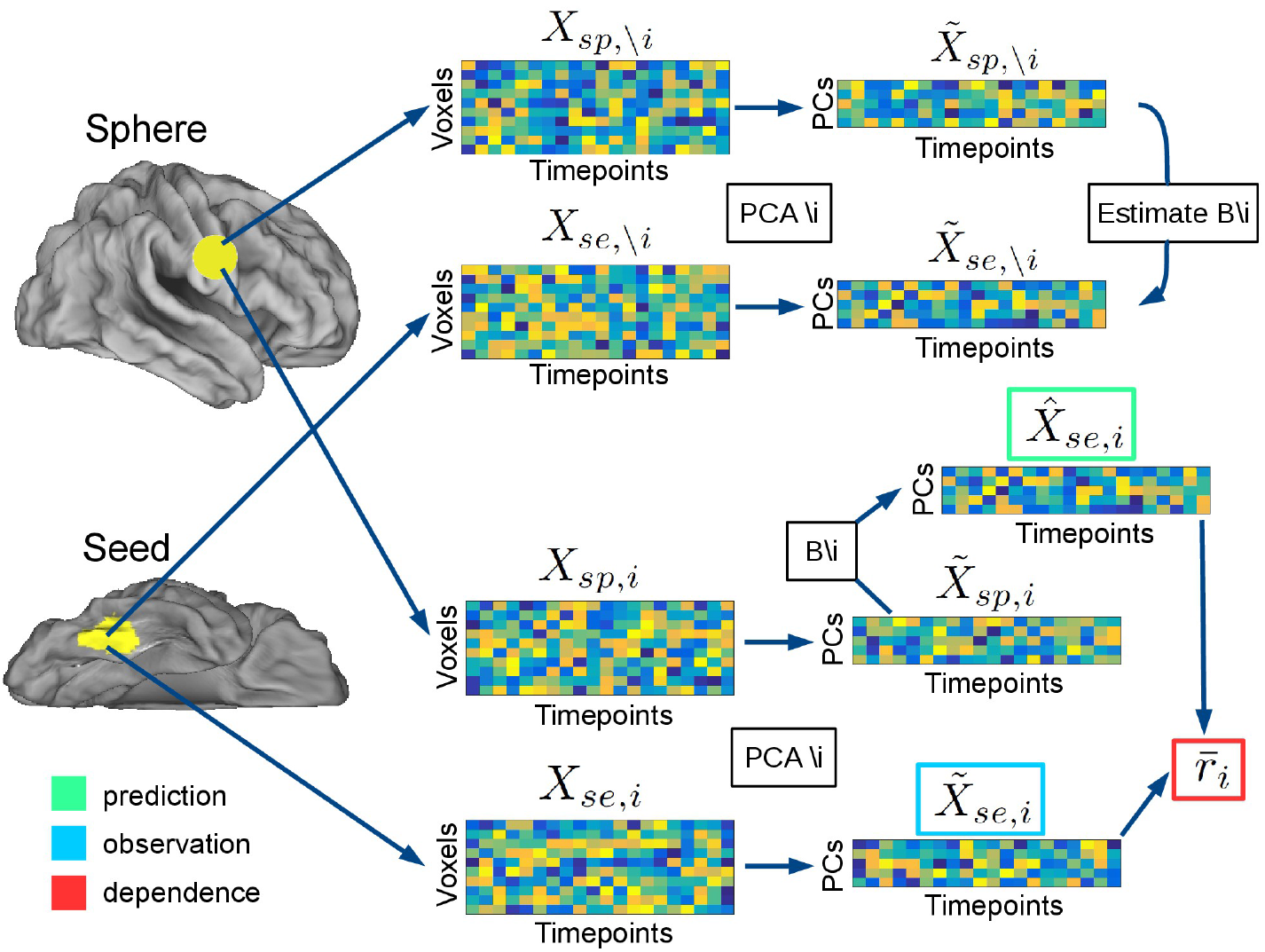
Analysis pipeline.

For each voxel *j* in the seed region, variance explained was calculated as

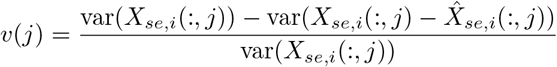

where 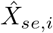 are the predicted voxelwise timecourses, and the values *v(j)* were averaged to obtain a single measure

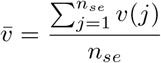

for each searchlight sphere.

## 3 Results

Standard functional connectivity identified statistical dependence between the FFA seed and other regions of ventral temporal cortex as well as with early visual cortex (peak MNI coordinates: 12,−90,−7), the right insula (peak MNI: 34,26,1), the thalamus (peak MNI: −9,−23,11), dorsal visual stream area V7 (14,−70,43) and intraparietal sulcus (IPS, peak MNI: 30,−66,32; Figure 3, in blue, FWE-corrected *p* < 0.05). MVPC additionally ientified statistical dependence between the FFA and the right superior temporal sulcus (rSTS, peak MNI: 51,−25,−4), the right anterior temporal lobe (rATL, peak MNI: 26,6,−33), right dorsomedial prefrontal cortex (rDMPFC, peak MNI: 8,57,30), posterior cingulate (pCing, peak MNI: 8,−46,38), and the dorsal visual stream area V3A (peak MNI 15,−88,31; Figure 3, in yellow, FWE-corrected *p* < 0.05). MVPC, unlike standard functional connectivity, did not detect significant statistical dependence between FFA and the amygdala (peak MNI for standard functional connectivity: 22,0,−20).

**Figure 3:**
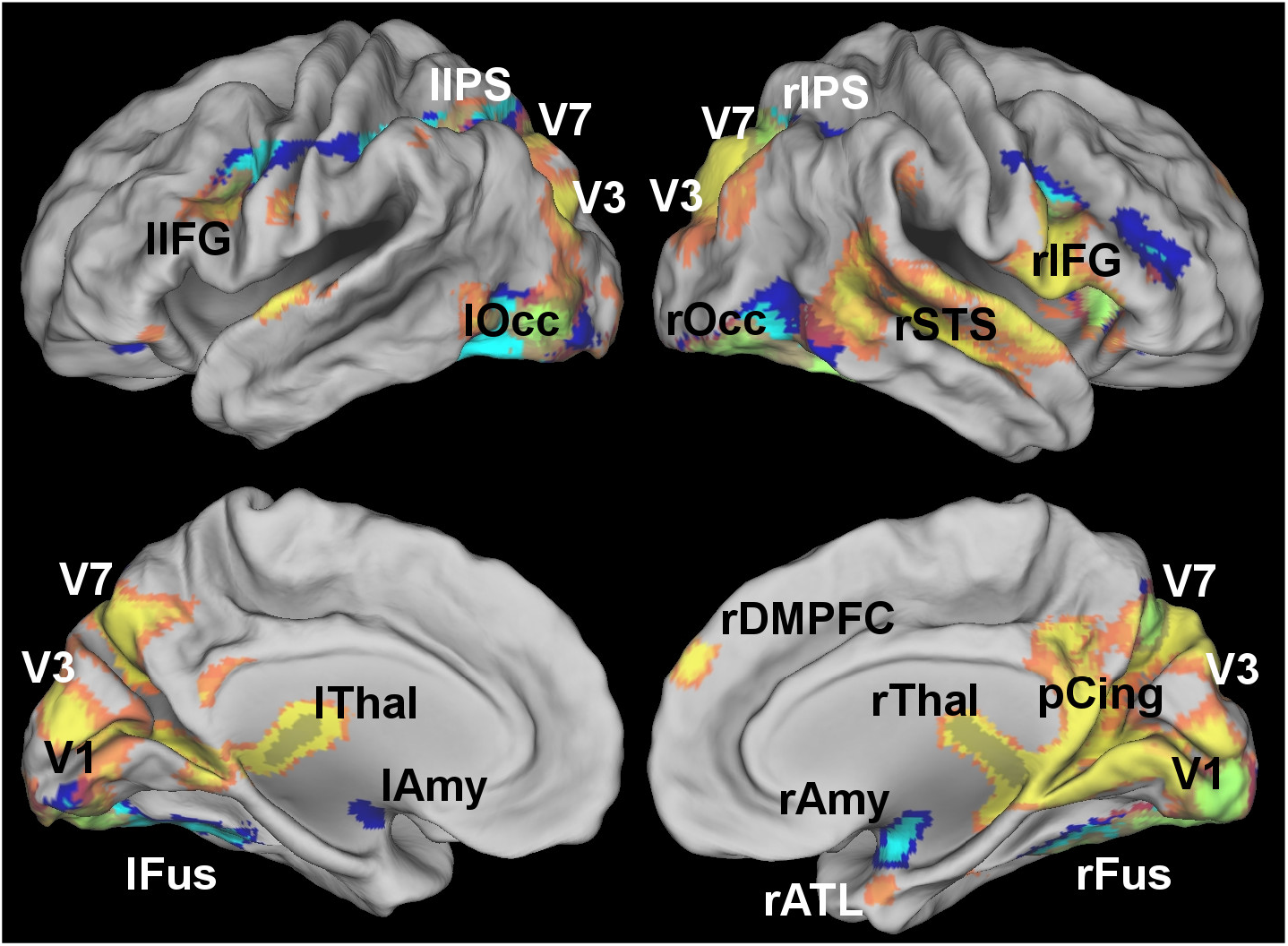
Brain regions showing statistical dependence with the FFA as identified by standard functional connectivity (blue) and multivariate pattern connectivity (MVPC, yellow) at a voxelwise FWE-corrected threshold *p* < 0.05. MVPC, but not standard functional connectivity, identified statistical depecn-dence with the right superior temporal sulcus, the right anterior temporal lobe, the right DMPFC and regions of the dorsal visual stream. Standard functional connectivity identified statistical dependence with the amygdala that was not detected by MVPC.

Analysis of voxelwise variance explained (see section 2.12) was performed for mean-based univariate connectivity, and for MVPC with 1, 2, and 3 principal components. Increasing the number of principal components led to a corresponding increase in the voxelwise variance explained in independent data (Figure 4 A for voxels explaining more than 5% of voxelwise variance, (Figure 4 B for voxels explaining more than 10% of voxelwise variance). As expected, the greatest amount of independent variance explained was observed in the right fusiform gyrus, in the proximity of the seed region’s location. Thanks to the additional contribution of the second and third principal components, variance explained above the 5% threshold was also observed more posteriorly extending towards the occipital face area (OFA), in the left fusiform, and anteriorly extending towards the medial portions of the anterior temporal lobes (ATL). These portions of cortex have been implicated together with FFA in the recognition of faces. (Nestor et al., 2011; Anzellotti et al., 2013; Anzellotti and Caramazza, 2015). The inclusion of dimensions beyond the first PC improved the modeling of statistical dependence between FFA and other regions implicated in face recognition. The voxelwise variance explained in independent data by mean-based univariate connectivity remained below 5% in the whole brain (Figure 4 C,D).

**Figure 4:**
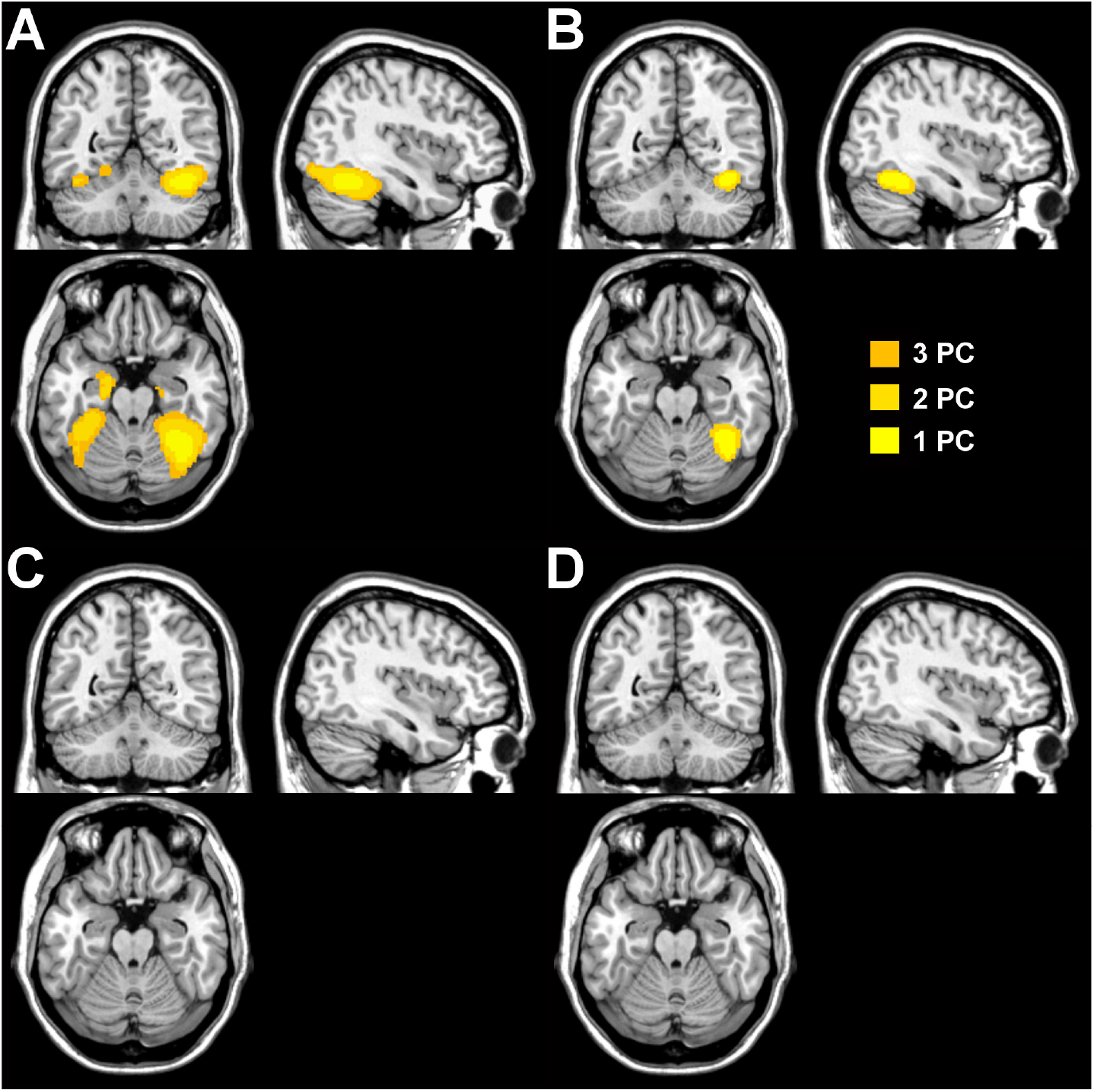
A) Regions explaining more than 5% of independent variance on average in FFA voxels, as measured by MVPC with 1, 2 and 3 principal components. B) Regions explaining more than 10% of independent variance on average in FFA voxels, as measured by MVPC with 1, 2 and 3 principal components. C) Regions explaining more than 5% of independent variance on average in FFA voxels, as measured by mean-based univariate connectivity. D) Regions explaining more than 10% of independent variance on average in FFA voxels, as measured by mean-based univariate connectivity.

Including additional dimensions beyond the first improved our ability to characterize the statistical dependence between responses in the FFA seed and responses in other brain regions that have been implicated in face processing. According to one hypothesis, each dimension of representational space in the FFA seed region could show statistical dependence with the same set of brain regions. In this case, the searchlight map for predicting the first PC in FFA would look like the searchlight map for predicting the second and the third PC. Alternatively, different dimensions of FFA’s representational space could show differential connectivity patterns with the rest of the brain. To investigate this question, we performed three separate MVPC-searchlight analyses, using only the first, second and third PC in FFA as seed respectively. The number of dimensions used in the spheres was determined with BIC. We then computed a similarity matrix between the resulting 27 searchlight maps for each participant and dimension using euclidean distance (Figure 5 A). The similarity matrix reveals that dimension 1 and dimensions 2,3 form two separate clusters. To test quantitatively this separation, we ran a permutation test subdividing the 27 searchlight maps in two random subsets and calculating average euclidean distance within vs between the random subsets. In none of 1000 permutation iterations, the between minus within distance was greater than in the subdivision assigning the maps obtained with the first PC to one subset, and the maps obtained with the second and third PCs to the other (*p* = 0.001). We then averaged the MVPC-searchlight maps for the first PC and for the second and third PCs, and we studied the spatial distribution of the top 5000 voxels in the brain showing greatest statistical dependence with the first PC (Figure 5, B in yellow) and the top 5000 voxels in the brain showing greatest statistical depen-dence with the second and third PCs (Figure 5, B in blue). The first PC showed greatest statistical dependence with voxels extending posteriorly towards early stages in the visual processing hierarchy, and dorsally towards regions in the dorsal visual stream. By contrast, the second and third PCs showed a different profile: strongest statistical dependence was found with regions extending anteriorly, towards the medial ATL. MVPC revealed different connectivity profiles for different dimensions of FFA’s representational space, individuating two subspaces showing disproportionate statistical dependence with regions involved in early and late visual processing respectively.

**Figure 5:**
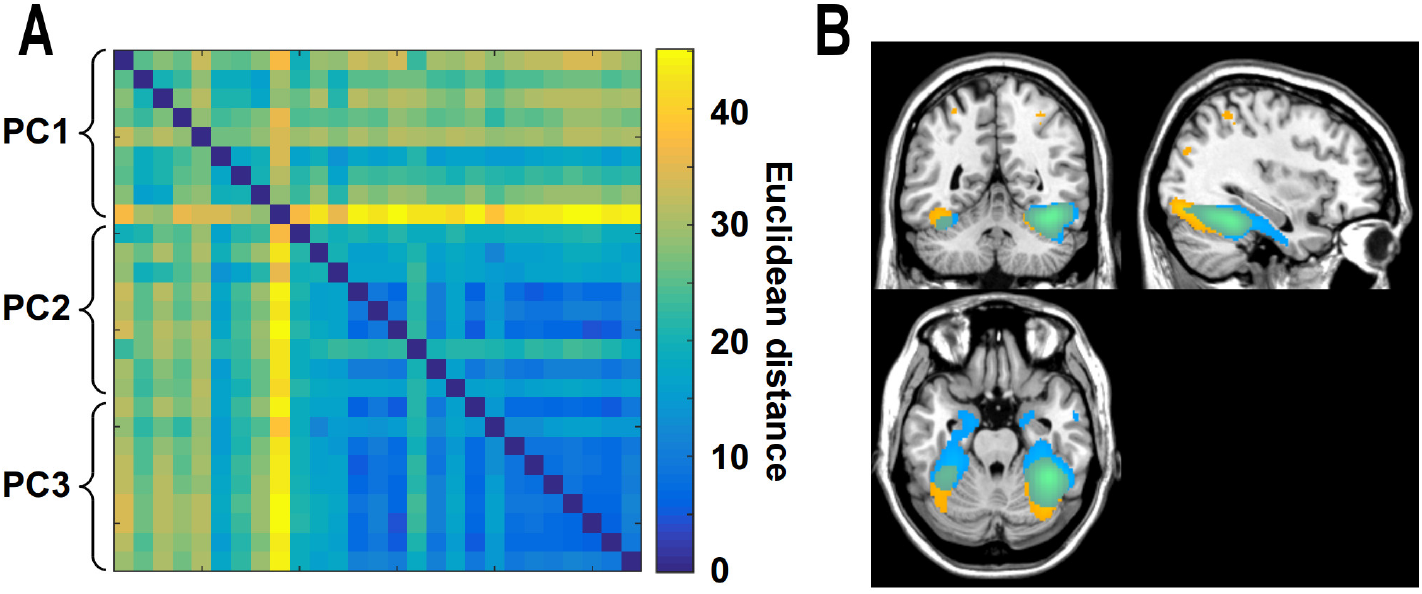
A) Similarity matrix between the whole-brain maps of 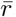 values obtained with MVPC for each participant reflecting the statistical dependence between each voxel and the first, second, and third PC respectively in the FFA seed. B) Top 5000 voxels showing highest 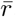 values for the first PC (in yellow), and for the second and third PCs (in blue). Different subspaces of FFA responses show different MVPC profiles, with the first dimension showing greatest statistical dependence with posterior ventral temporal regions and regions in the dorsal visual stream, and the second and third dimensions showing greatest statistical dependence with anterior temporal regions.

## 4 Discussion

This article introduces multivariate pattern connectivity (MVPC), a new method to investigate the multivariate statistical dependence between brain regions. MVPC is characterized by several key properties. First, the BOLD signal in each brain region is modeled as a set of responses along multiple dimensions, with each dimension corresponding to a function of the voxels in that region. Second, MVPC investigates the statistical dependence between two regions by computing the extent to which the responses in the multiple dimensions characterizing one region can predict the responses in the multiple dimensions characterizing the other region over time. Third, with an analogy to MVPA methods, MVPC uses a cross-validation procedure in which independent data are used for training and testing of the models. A subset of the runs are used as a training set to generate parameters which are then tested assessing their ability to predict responses in a left-out independent run. This leave-one-out approach mitigates the impact of noise, improving on most current methods that do not test the extent to which relationships between regions are sufficiently stable to generalize to independent data.

In the examples described in the present article, dimensions are obtained with PCA as linear combinations of the voxels that tend to be jointly activated or deactivated over time. From a neuroscientific perspective, we can think of each region as consisting of multiple neural populations with selectivities for different properties of the stimuli that have different distributions over the course of the experiment. Each population has different spatial distributions over voxels. This leads different weighted combinations of voxels to having different timecourses of responses, whose dynamics can provide deeper insights into the interactions between regions than the investigation of average responses. Of course,while different populations withdifferent selectivities and different spatial distributions can lead to dimensionswithdifferent time courses, it is unlikely thatindividual dimensions obtainedwith PCA correspond in a one-to-one relationship to neural populations witha specific selectivity profile. For example, more than one neural population might be collapsed in a single principal component, or populations might not be assigned to dimensions in a one-to-one mapping because of the orthogonality constraints imposed by PCA.

Like standard functional connectivity, MVPC revealed statistical dependence between the FFA and more posterior regions of ventral temporal and occipital cortex, and with regions in early visual cortex. However, MVPC additionally revealed statistical dependence between the FFA and the right ATL, previously implicated in the recognition of face identity (Kriegeskorte et al., 2007; Nestor et al., 2011; Anzellotti et al., 2013; Anzellotti and Caramazza, 2015). Furthermore, MVPC (but not standard functional connectivity) identified statistical dependence between the FFA and the right STS, implicated in the recognition of person identity (Winston et al., 2004; Anzellotti, 2014; Hasan et al., 2016; Anzellotti and Caramazza, 2017) and of facial expressions (Peelen et al., 2010; Skerry and Saxe, 2014; Deen et al., 2015). Standard functional connectivity, but not MVPC, identified statistical dependence between FFA and the amygdala. As was briefly mentioned in section 2.12, this can be due to less stable predictive relationships between responses in the amygdala and FFA dimensions beyond the first PC.

MVPC led to important improvements in independent variance explained at the voxel level over mean-based univariate connectivity (Figure 4). MVPC using a single principal component already improved variance explained over a mean-based univariate approach. Adding a second and a third PC further improved variance explained in ventral temporal cortex as well as the anterior temporal lobes. In the end, MVPC allowed us to separately investigate the connectivity profiles of different dimensions of FFA’s representational space.

MVPC differs in important respects from previous techniques aimed at studying the dynamic interactions between brain regions in terms of the information they encode. Unlike previous techniques (Coutanche and Thompson-Schill, 2014; Henriksson et al., 2015), MVPC does not rely on discrimination between categories determined by the experimenter, but on dimensions derived in a data-driven fashion. The data-driven dimensions can be related to properties of the stimuli or the task with a subsequent model (for instance regressing dimensions on conditions, or on stimulus properties using a forward model). Another difference between MVPC and the methods introduced by Coutanche and Thompson-Schill (Coutanche and Thompson-Schill, 2013, 2014) is that the latter characterize each region with a single measure (how well the pattern in a given timepoint can be assigned to one condition or another), while MVPC adopts multiple measures (the values along the multiple dimensions), which can provide a richer characterization of a region’s representation at any given time. An innovative study (Henriksson et al., 2015) investigated the relations between brain regions measuring the correlation between representational dissimilarity matrices in different regions. This approach provides a richer characterization of each region’s representational structure by comparing similarity matrices instead of classification accuracies, but it discards trial-by-trial variability. Furthermore, correlations between dissimilarity matrices can only be computed if the same set of conditions are used to generate the dissimilarity matrices in each region. When the conditions correspond to individual stimuli as in Henriksson et al. (2015) this is not problematic, but if stimulus categories are used it raises the question of whether it is appropriate to characterize the representational spaces of very different brain regions in terms of the dissimilarities between the same set of categories. Taking images of objects as an example of stimuli, categorization based on animacy could be most appropriate for some brain regions, while categorization based on color could be more appropriate for other regions. An additional approach has used distance correlation to capture multivariate dependences between regions Geerligs et al. (2016), finding more robust results than traditional correlations for inhomogeneous regions. MVPC offers as advantages over this approach the ability to test stability of the dependence between two regions in independent data, and to analyze dependence for different representational subspaces (e.g. Figure 5). This feature of MVPC makes it possible to relate the dimensions characterizing a region’s responses to stimulus properties using forward models, to then investigate what representational content drives statistical dependence between two regions.

However, the most important asset of MVPC is probably its flexibility. The framework of 1) modelling representational spaces in individual regions, 2) considering multivariate timecourses as trajectories in these representational spaces, and 3) fitting models predicting the trajectory in the representational space of one region as a function of the trajectory in the representational space in another offers a wealth of possibilities to build increasingly refined models, both in terms of the characterization of representational spaces and in terms of the models of their interactions. For the characterization of representational spaces, in this article we adopted PCA as a simple example, but other methods such as independent component analysis (ICA) and nonlinear dimensionality reduction techniques can also be used. For modelling interactions between regions, we limited the current application to simultaneous, non-directed interactions following an approach similar to functional connectivity, but MVPC makes it possible to model nonlinear maps between representational spaces (Anzellotti et al., 2016), and to use models that investigate the directionality of interactions using temporal precedence, along the lines of Granger Causality (Roebroeck et al., 2005), Dynamic Causal Modelling (Friston et al., 2003), and Dynamic Network Modelling (Anzellotti et al., 2017).

## Acknowledgments

This study was supported by the Center for Mind/Brain Sciences (CIMeC) of the University of Trento and by NIH Grant 1R01 MH096914-01A1 to Prof. Rebecca Saxe. Stefano Anzellotti was supported by a Postdoctoral Fellowship from the Simons Center for the Social Brain.

